# scSAID: A Comprehensive Cross-Species Single-Cell Skin Atlas Reveals Species-Specific Responses to Psoriasis

**DOI:** 10.64898/2026.07.20.739409

**Authors:** Yixiang Ren, Yuchen Shen, Linxi Jin, Yufei Huang, Yueshan Deng, Ying Xiao, Chaochen Wang

## Abstract

Single-cell RNA sequencing has rapidly expanded the scale of skin transcriptomic data, yet these datasets remain fragmented across studies spanning different species, diseases and experimental manipulations. An up-to-date, comprehensive, and queryable single-cell cross-species repository for skin is still lacking. Here, we present <u>scSAID</u> (skin-scsaid.com), a single-cell database with an interactive web portal offering a broad suite of in-depth analyses for human and mouse skin. It integrates more than 1.2 million high-quality cells collected from 252 samples, establishing a unified reference for cell-type annotation, cross-species comparison and pathological studies. Using psoriasis as a case study, we demonstrate how scSAID can be used to evaluate how faithfully mouse models reproduce human pathology. Systematic comparison with the imiquimod-induced mouse model revealed numerous species-specific molecular signatures of psoriasis, including human-specific *NFKB1* activation and *STAT1* involvement, indicating that the current mouse model captures only limited aspects of the disease. We further introduce psoSpotter, a disease-biomarker-selection algorithm that, coupled with in silico perturbation using scSAID data, uncovers *PPIA* as a novel psoriasis drug target, illustrating the potential of scSAID for identifying therapeutic approaches. Overall, scSAID delivers a large-scale, cross-species skin single-cell resource and analysis platform, opening new opportunities for the discovery of disease-relevant targets in skin diseases.

## 1 Introduction

The skin is a major barrier organ that protects the body from environmental injury and infection^1^. It contains diverse epithelial, immune, stromal, vascular and neural populations that together maintain tissue homeostasis and respond to external hazards^2,3^. Skin and subcutaneous diseases impose a substantial global burden, particularly through disability and reduced quality of life^4^ and impose substantial clinical, psychological and socioeconomic burdens^5^. A 2020 modelling study estimated self-reported lifetime psoriasis prevalence in adults at up to 1.12% globally^6^. Understanding the cellular composition of skin and how its cellular and molecular signature is altered in disease is therefore essential for elucidating disease mechanisms and informing mechanistic and translational research.

Single-cell RNA sequencing (scRNA-seq) has transformed the study of skin biology by enabling cell-type-resolved profiling of skin. An increasing number of studies have applied single-cell transcriptomics to inflammatory skin diseases, malignancies and wound healing^7,8^, as well as to cutaneous infections such as leprosy ^9,10^, collectively generating a valuable public resource for investigating skin cell states and disease-associated molecular profiles. However, leveraging this resource at scale remains challenging because the datasets are distributed across independent repositories and differ in sampling strategy, metadata structure, tissue processing and cell-type nomenclature^3^. Study-specific choices in quality control, normalization, dimensionality reduction, batch correction and annotation further complicate direct comparison across datasets.

To address this need, we developed <u>scSAID</u> (<u>s</u>ingle-<u>c</u>ell <u>s</u>kin and <u>a</u>ppendages integrated <u>d</u>atabase), an integrative cross-species skin scRNA-seq database and public interactive web platform, freely accessible at skin-scsaid.com. scSAID aggregates 405,484 human cells from 133 samples and 794,766 mouse cells from 119 samples, spanning 18 human and 22 mouse conditions, built entirely from raw sequencing data with a customized processing pipeline. The platform supports both single-dataset-level and atlas-level exploration through an array of in-depth analysis functions such as cell–cell communication inference, regulatory analysis and cross-dataset gene-centered queries. It further links analytical outputs to the original study context for large language model (LLM)-assisted interpretation, generating context-aware summaries of the corresponding results and suggesting new insights, making it suitable for both experts and new users.

To illustrate how scSAID turns integrated data into biological insight, we focus on psoriasis, a common immune-mediated skin disease driven by a self-amplifying IL-23/IL-17 axis that couples keratinocyte hyperproliferation with sustained T-cell-driven inflammation^6^. Although the IMQ model reproduces several inflammatory and histopathological features of human psoriasis, its correspondence to the original human psoriasis at single cell resolution remains incompletely defined^11,12^.Using scSAID, we compared human and mouse psoriasis to define conserved and species-specific features of the imiquimod model. We further developed psoSpotter, a disease-agnostic biomarker-identification algorithm that selects a minimal, sex- and age-informed gene panel with potential clinical utility, and applied it to characterize cross-species molecular footprints in psoriasis. Coupled with in silico perturbation and drug-database resources, we nominated PPIA as a candidate psoriasis target for experimental validation.

## 2 Results

### 2.1 Overview of scSAID workflow and website functions

scSAID is built through dataset aggregation, species-specific preprocessing and integration, and web-based deployment. Public-domain 10x skin scRNA-seq datasets were collected from two omics databanks, the Gene Expression Omnibus (GEO) of National Center for Biotechnology Information (NCBI) and the Genome Sequence Archive (GSA) of National Genome Data Center (NGDC), and organized into human and mouse cohorts covering healthy, diseased and experimentally perturbed skin states. The human cohort comprised 405,484 high-quality cells from 133 samples across 23 GEO or GSA accessions, spanning healthy donors and 17 disease conditions, including psoriasis, hidradenitis suppurativa, lymphoma, actinic keratosis, melanoma and squamous cell carcinoma. The mouse cohort comprised 794,766 cells from 119 samples across 26 GEO or GSA accessions, including 64 healthy samples and 21 perturbation or disease conditions, such as the imiquimod (IMQ)-induced psoriasis-like model, mechanical wounding, genetically induced carcinoma and multiple engineered genetic backgrounds **(Figure 1a)**. In total, scSAID integrates 1,200,250 skin and appendage cells from 252 samples across 49 studies, providing a cross-species reference across the sampled conditions and sites **(Figure 1a)**.

**Figure 1.**
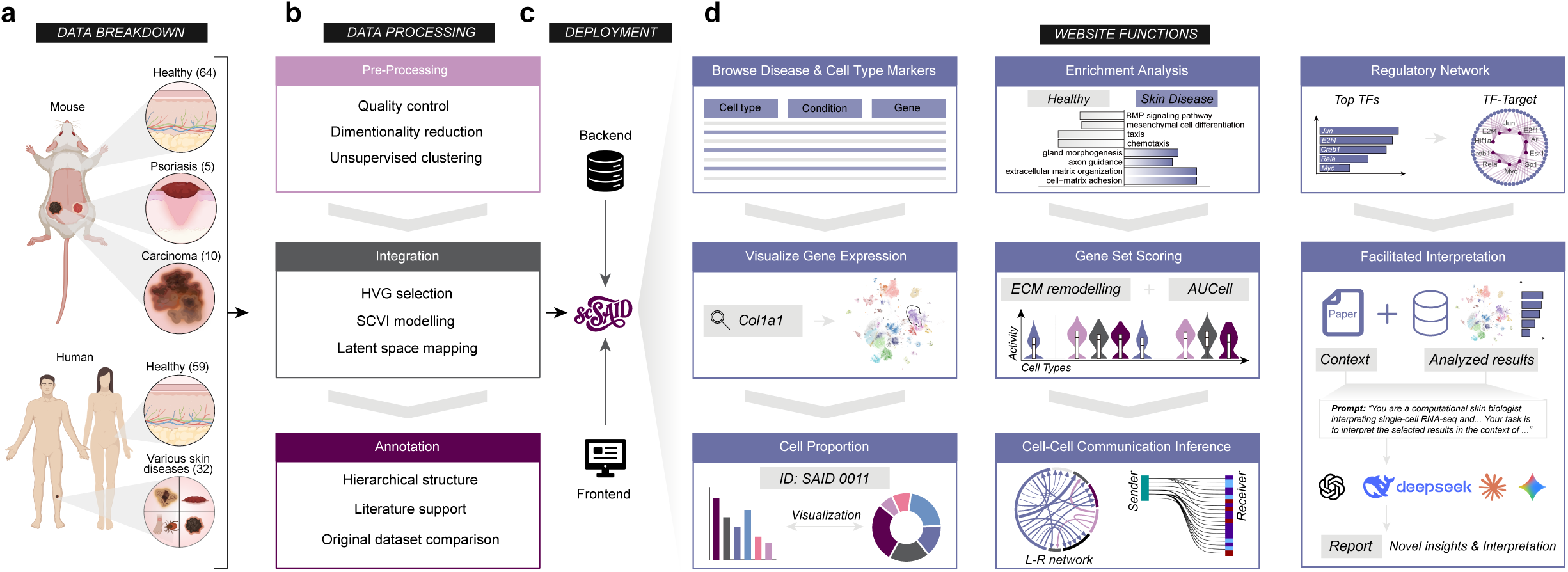
scSAID overview. **(a)** Dataset breakdown for mouse and human, stratified by broad condition category. **(b)** Data-processing pipeline of scSAID. **(c)** Backend–frontend deployment of the website portal. **(d)** Eight selected core analysis functions available through the scSAID website.

To ensure that datasets generated across independent studies could be compared within a unified framework, all samples were processed using a standardized, within-species preprocessing workflow. Raw sequencing data were processed independently at the sample level to remove low-quality cells, technical artifacts and doublets, while preserving raw counts for downstream analysis. Per-sample dimensionality reduction, clustering and uniform manifold approximation and projection (UMAP) visualization were examined one by one to assess data quality and sample-level structure prior to integration. A similar preprocessing strategy was applied to human and mouse datasets, with parameters tailored to each species (see Methods for details).

Human and mouse samples were then integrated separately to generate species-specific reference atlases **(Figure 1b)**. After excluding very small batches, highly variable genes were reselected from raw counts, and scVI^13^ was trained to learn a batch-aware latent representation while accounting for study-level and quality-related covariates. The resulting latent spaces supported integrated neighbor-graph construction and subsequent resolution-based clustering. Visual inspection of study and batch distributions on the UMAP was used to assess whether cells from independent accessions were embedded within shared biological compartments. After integration, cell identities were assigned by combining differential-expression analysis to identify cluster-specific markers with annotation based on canonical markers (**Supplementary Figures S1,S2**; Supplementary Table S1). Both the human and mouse atlases were annotated at two levels: broad major cell types and refined subtypes identified through lineage-specific subclustering.

The final atlas objects were deployed as an interactive public web resource designed to make the integrated datasets directly reusable **(Figure 1c)**. The scSAID web portal provides the integrated human and mouse objects separately for custom exploration, together with a suite of in-depth, browser-based analysis tools. Users can browse datasets by species, condition, tissue site, and study, and examine cell-type-specific and disease markers across datasets. Within a given dataset, users can visualize expression of genes of interest; inspect cell proportions and clustering structure; and perform differential-expression analysis, enrichment using gene set enrichment analysis (GSEA) or over-representation analysis (ORA) with GSEApy^14^, gene-set scoring with AUCell^15^, UCell^16^, single-sample GSEA (ssGSEA) ^17^, or gene set variation analysis (GSVA) ^18^, cell–cell communication inference and transcription-factor regulatory-network analysis^19,20^. Selected outputs can additionally be passed to an LLM-assisted interpretation function, supporting ChatGPT, Claude, DeepSeek, and Gemini with retrieval-augmented generation (RAG) capability, which contextualizes results against the original publication and generates hypotheses through a curated prompt **(Figure 1d)**. Together, these functions make the atlas queryable across exploring skin cell states, disease-associated remodeling, and candidate regulatory mechanisms.

### 2.2 Overview of scSAID data

scSAID spans multiple biological and anatomical categories in both mouse and human. The mouse atlas spans embryonic to aged postnatal stages from E13.5 to P700 and includes 22 condition categories across 11 anatomical-site categories, covering major skin and appendage-related regions such as dorsal skin, ear, footpad, tail, whole trunk skin, and back skin **(Figure 2a)**. The human atlas spans donor ages from 15 to 76 years, with an average age of 45 years, and includes 18 condition categories across nine anatomical-site categories, including whole skin, scalp, subcutaneous tissue, ear, foreskin, and Skoog’s fascia **(Figure 2b)**. In addition, scSAID is built from datasets with heterogeneous but broadly comparable sampling depths. The median number of cells per sample was 5,940 for mouse and 2,011 for human, with most samples contributing between 1,000 and 10,000 cells **(Figure 2a,b)**. Across 16 previously published skin-focused or whole-body single-cell atlas resources containing skin tissue^3,21–35^, scSAID contained the largest number of skin cells among the skin-dedicated resources included in our comparison, comprising approximately 1.2 million cells across the human and mouse atlases **(Figure 2c)**.

**Figure 2.**
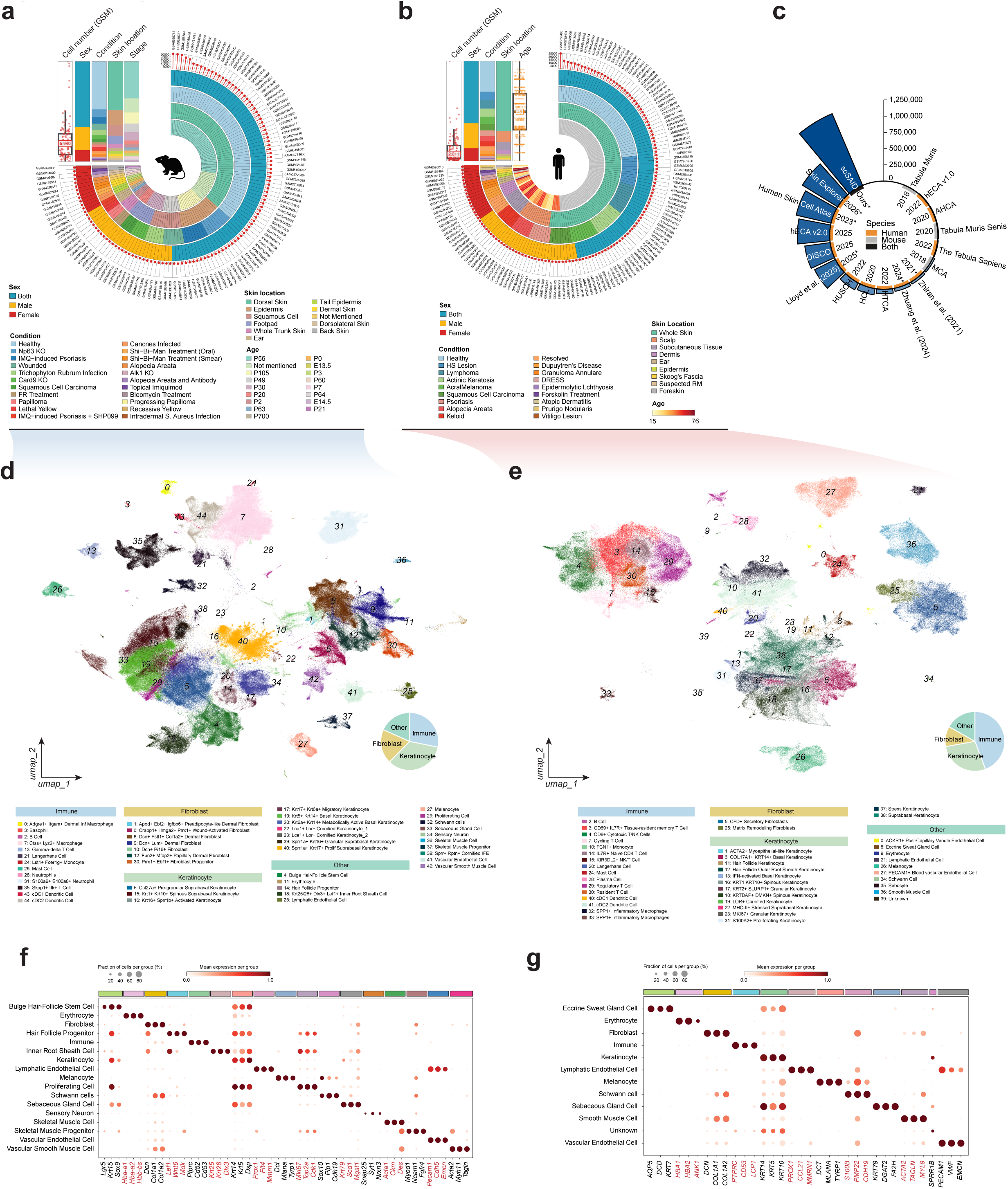
Mouse and human atlas breakdown and comparison. **(a,b)** Metadata manifest for the mouse **(a)** and human **(b)** cohorts, showing per-GSM cell number, sex, condition, skin location, and developmental stage or donor age. **(c)** Cell-number comparison of scSAID against 16 previously published skin or whole-body single-cell atlases that contain skin tissue. The year of publication for each atlas is marked on the inner circle at the base of each bar. Skin-dedicated resources are indicated with an asterisk after the year. **(d,e)** scVI-integrated UMAP embedding for the mouse **(d)** and human **(e)**, with broad-lineage proportions shown as inset pie charts. **(f,g)** Dot plots showing canonical lineage markers for the mouse **(f)** and human **(g)** atlas cell-type annotations, supporting the annotations shown in **(d,e)**.

Species-level integration resolved the cellular architecture of human and mouse skin into broad lineage classes and refined cell-state annotations. The mouse atlas contained 45 fine-grained cell types, whereas the human atlas contained 41 fine-grained cell types **(Figure 2d,e)**. In both species, the integrated atlases recovered the major skin-resident compartments, including epithelial, immune, stromal, vascular, melanocytic, and glandular cells. The mouse atlas additionally captured hair-follicle and inner-root-sheath compartments, whereas the human atlas included eccrine sweat gland populations and other human skin-specific compartments. Cells were annotated using markers from validated, previously published studies for both human and mouse (**Figure 2f,g**; Supplementary Table S1), as was the finer subdivision within each atlas **(Supplementary Figures S1 and S2)**. In each case, marker expression was restricted to the expected populations, supporting the accuracy of the annotation.

As a further check on annotation reliability, we compared scSAID cell-type assignments with source-study expectations in three randomly selected datasets **(Supplementary Figure S3)**. Human acral melanoma samples from GSE242477, which profiled tumor-infiltrating lymphocytes isolated by fluorescence-activated cell sorting (FACS) for CD3^+^CD45^+^ cells, were assigned almost entirely to the immune compartment. Isolated mouse epidermis from the *K14-CreT2* ΔNp63 model (GSE213528) showed the expected keratinocyte enrichment. Full-thickness back-skin samples from the DMBA/TPA-driven multistage cutaneous squamous-cell carcinoma series (GSE261766) retained the expected stage-associated compositional pat-terns^36–38^. These concordances support the accuracy of cell-type annotation.

### 2.3 scSAID reproduces the type 17 T-cell signature in recurrent psoriatic plaques

To demonstrate the integrated analysis workflow implemented in scSAID and its utility, we reanalyzed the psoriasis dataset GSE248121 selected through the dataset-browsing interface **(Figure 3a)**. The source study profiled on-site recurrent psoriatic plaques, previously resolved lesions and clinically normal-appearing skin from four patients. The original study identified IL-17A/IL-17F double-producing T cells, phenotypically unstable regulatory T cells and quiescent tissue-resident memory T cells as candidate populations associated with local recur-rence^39^. We therefore asked whether scSAID could recapitulate these T-cell states, relate them to keratinocyte responses and predicted intercellular and regulatory programs, and generate hypotheses not reported in the source study.

**Figure 3.**
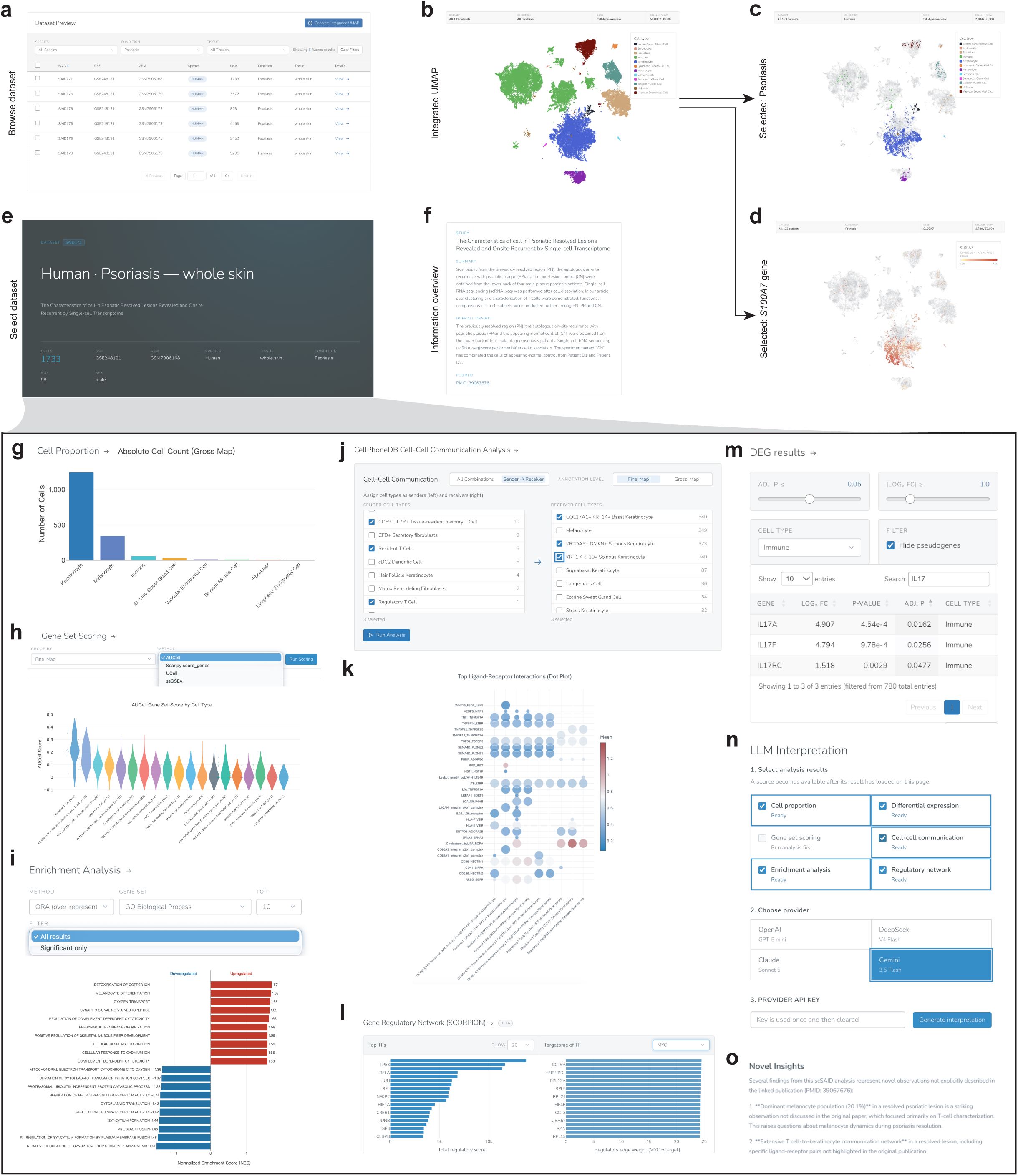
Demonstration of scSAID website portal functionality on SAID171. **(a)** Interface for dataset browsing, from which SAID171 (GSE248121) was selected. **(b–d)** Integrated human skin atlas visualized by UMAP and colored by broad cell type **(b)**, followed by restriction to cells from psoriasis samples **(c)** and visualization of *S100A7* expression within the selected condition **(d)**. **(e,f)** SAID171 dataset detail page **(e)** showing sample metadata and the linked study overview **(f)**, including the study summary, experimental design and publication source. **(g)** Absolute cell counts across broad cell-type annotations of SAID171. **(h)** Gene-set scoring interface showing AUCell-based activity across fine-grained cell types. **(i)** GSEA showing positively and negatively enriched GO biological processes. **(j)** Cell–cell communication inference interface for selecting sender and receiver populations. **(k)** Dot-plot representation of the highest-ranked ligand–receptor interactions across selected sender–receiver pairs. **(l)** Gene-regulatory-network analysis showing the highest-scoring TFs and the inferred targetome of a selected regulator, exemplified by *MYC*. **(m)** Interactive DEG table filtered for immune cells. **(n)** LLM-interpretation panel for selecting scSAID-derived results coupled with original study context. **(o)** Example snapshot of LLM-generated insights on the website (truncated).

The integrated UMAP resolved the major cellular compartments represented in the atlas **(Figure 3b)**. After restricting the atlas to psoriasis datasets, cells were concentrated predominantly within keratinocyte clusters **(Figure 3c)**. We therefore examined *S100A7*, a keratinocyte alarmin induced by cytokines produced by type 17 T cells^40^. Consistent with its epidermal origin, *S100A7* expression was concentrated within keratinocyte clusters **(Figure 3d)**. We next selected SAID171 (GSM7906168) for analysis at single-sample level **(Figure 3e,f)**. The sample contained 1,733 cells, of which 1,248 were keratinocytes, accounting for 72.0% of the total population **(Figure 3g)**.

To determine whether SAID171 retained the inflammatory T-cell states emphasized by the source study, we used AUCell to score a literature-derived type 17 T-cell gene set^41–45^. Resident T cells showed the highest score distribution, followed by CD69^+^IL7R^+^ tissue-resident memory T cells **(Figure 3h)**. This pattern was consistent with the source study’s emphasis on tissue-resident T-cell populations^39^.

We compared SAID171 with the remaining datasets in the atlas to place these cell-state results in a broader transcriptional context and performed Gene Ontology (GO) enrichment analysis on the resulting differentially expressed genes (DEGs). Genes increased in SAID171 were enriched for complement-dependent cytotoxicity and responses to metal-ion stress, whereas decreased genes were associated with cytoplasmic translation and mitochondrial electron transport **(Figure 3i)**. These enrichments defined a broader dataset-associated inflammatory and metabolic context for the T-cell states identified above.

We next inferred candidate T cell–keratinocyte ligand–receptor interactions in SAID171 using the cell–cell communication inference function on our website via CellPhoneDB v5 **(Figure 3j,k)**. Predictions involved WNT, semaphorin and TNF signaling between resident or regulatory T cells and basal or spinous keratinocytes. These included WNT16–FZD6/LRP5 and TNF–TNFRSF1A interactions involving resident T cells, together with SEMA4D–PLXNB1 and SEMA4D–PLXNB2 interactions involving CD69^+^IL7R^+^ tissue-resident memory T cells. Notably, CellPhoneDB predicted IL-17F- and IL-17A/F-mediated signaling from resident T cells to KRTDAP^+^DMKN^+^ spinous keratinocytes through the IL-17RA/IL-17RC receptor complex (Supplementary Table S2). These predictions linked resident T-cell populations with high type 17 signature scores to a defined spinous keratinocyte population. To assess whether these communication patterns were reproducible in another psoriasis sample, we repeated the analysis in SAID173. Most of the detected ligand–receptor interactions were shared between SAID171 and SAID173, including interactions within the IL-17 axis (Supplementary Table S2). Gene regulatory network (GRN) analysis subsequently identified the transcription factors (TFs) *TP53*, NF-*κ*B-family and AP-1-family among the prominent candidate regulators in SAID171 **(Figure 3l)**. The GRN analysis extended the cell-state and communication results by placing them within an inferred upstream regulatory context that was not determined by the original study.

We also tested whether the sample-level type 17 signal was detectable across the atlas by performing pseudobulk differential expression analysis of immune cells from psoriasis and healthy skin. Immune cells from psoriasis samples showed increased expression of *IL17A* (adjusted *P* = 0.016) and *IL17F* (adjusted *P* = 0.026). *IL17RC* was also modestly increased (adjusted *P* = 0.048; **Figure 3m**). Thus, complementary analyses of overlapping data, including gene set scoring at the sample level, communication inference, and pseudobulk differential expression across the atlas, consistently indicated type 17 immune activity in psoriasis, in agreement with the central finding of the source study^39^.

Finally, the LLM-assisted interpretation module allowed us to jointly select and interpret outputs from cell-composition, differential-expression, cell–cell communication, enrichment and regulatory-network analyses **(Figure 3n)**, together with the context provided by the original paper. In this example, we used Gemini 3.5 Flash to generate a single natural-language report from the selected outputs **(Figure 3o; Supplementary Figure S4)**. In combination, these scSAID functions reproduced selected findings of the source study and generated additional, testable hypotheses.

### 2.4 Cross-species analysis reveals limited conservation of immune cells in the mouse IMQ model

To assess how reliably the IMQ-induced mouse model reproduces molecular signatures of human psoriasis, we focused on the immune compartment, which has a central role in psoriasis pathogenesis and treatment^6^. We therefore performed an integrated cross-species analysis using human and mouse whole-skin psoriasis datasets obtained from scSAID.

Perturbation scoring with Augur^46^ identified substantial disease-associated perturbation of immune cells in both species **(Figure 4a,b)**. Pseudobulk differential expression analysis was performed within the immune compartment between healthy and psoriasis conditions in the two species. After ortholog mapping, 500 genes were specific to the mouse comparison, 853 were specific to the human comparison, and 25 were significant in both species (**Figure 4c**; Supplementary Table S3). The cross-species comparison therefore comprised 525 mouse and 878 human disease-associated genes and revealed limited overlap between the two immune transcriptional responses.

**Figure 4.**
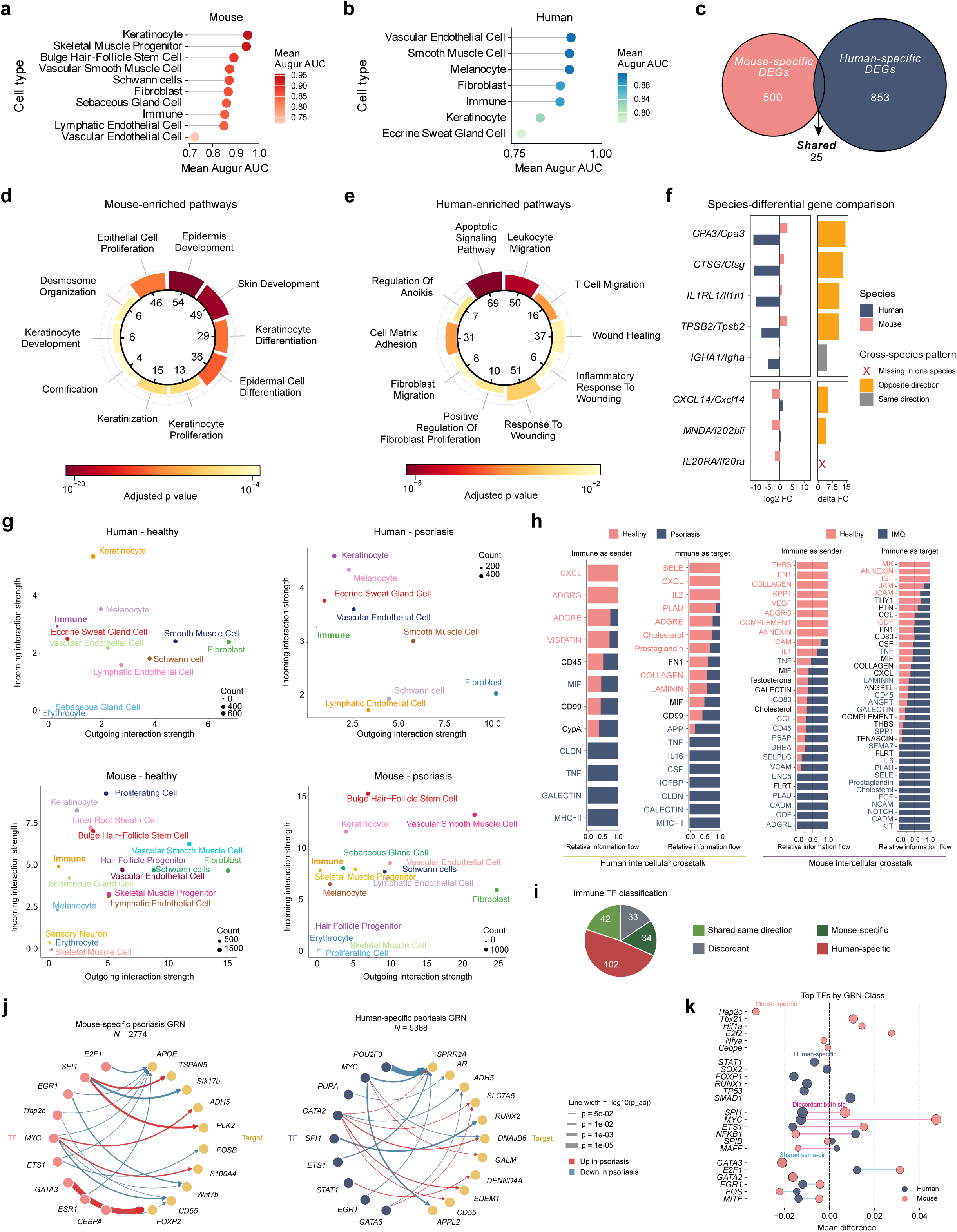
Cross-species comparison of the immune compartment in psoriasis. **(a,b)** Augur perturbation AUC per cell type for the mouse **(a)** and human **(b)** cohorts, comparing healthy versus IMQ-induced psoriasis conditions. **(c)** Venn diagram of immune-compartment DEGs after one-to-one human-ortholog mapping. **(d,e)** Top GO biological-process terms enriched among mouse-specific **(d)** and human-specific **(e)** immune DEGs, with adjusted *P* value indicated by color and enriched gene count by inner ring sector. **(f)** Representative species-differential genes in immune cells shown as paired log_2_ fold change in human and mouse (left) and as cross-species delta FC (right). **(g)** Cell-type-level outgoing versus incoming interaction strength calculated by CellChat for human and mouse, separately for healthy and psoriasis or IMQ contexts. **(h)** Pathway-level information flow calculated by CellChat for the immune compartment as either signaling sender or signaling target, comparing healthy versus disease in each species. Significant pathways are shown in color. **(i)** Cross-species classification of significant immune-compartment TFs from GRN inference. **(j)** Mouse-specific (left) and human-specific (right) psoriasis GRN subnetworks, showing edges that differ significantly between the disease and healthy networks within each species, with line width mapped to *P* value and color mapped to direction of regulation. **(k)** Top TFs stratified by GRN class, plotted as the disease-versus-healthy mean difference in TF activity for the mouse and human cohorts.

Enrichment analysis of the species-specific gene sets identified distinct biological programs. Human-specific genes were enriched for leukocyte and T-cell migration, wound healing, and cell–matrix adhesion, whereas mouse-specific genes were enriched for epidermal development, keratinocyte differentiation, and keratinization **(Figure 4d,e)**. Inspection of the genes contributing to the mouse-specific enrichment showed that these terms were driven largely by junction- and adhesion-associated genes, including *Cdh3*, *Dsg3*, and *Itgb6* **(Figure 4d,e)**. By contrast, *Krt* and *Sprr* genes included in the study’s keratinocyte-marker set accounted for only approximately 4% of the mouse-specific gene set and were predominantly downregulated following IMQ treatment. Thus, the mouse-specific enrichment reflected a restricted junctional and epithelial-remodeling-associated gene program. Directionally discordant responses were also apparent at the individual-gene level **(Figure 4f)**. The mast-cell-associated genes *CPA3*, *TPSB2*, and *IL1RL1*, together with the serine protease gene *CTSG*, decreased in human psoriasis but increased following IMQ treatment in mouse skin. *CXCL14* and the *MNDA/Ifi202b* ortholog pair showed a similar opposite-direction pattern. By contrast, *IGHA1/Igha* changed in the same direction in both species, whereas *IL20RA/Il20ra* was only captured in mouse **(Figure 4f)**.

Cell–cell communication inference further identified both conserved and divergent disease-associated changes. The immune compartment showed increased outgoing interaction strength in human psoriasis and in IMQ-treated mouse skin **(Figure 4g)**. However, pathway-level relative information flow differed extensively between species (**Figure 4h**; Supplementary Table S4). TNF signaling showed increased disease-associated information flow in both comparisons. In contrast, inferred E-selectin (SELE) information flow was higher in healthy human skin but in IMQ-treated mouse skin, indicating an opposite pattern between the human disease and mouse model. Notably, a previous randomized trial of the anti-E-selectin antibody CDP850 reduced E-selectin abundance but did not significantly improve PASI scores in patients with chronic plaque psoriasis ^47^.

GRNs (Gene Regulatory Networks) inferred using SCORPION^20^ classified 211 identified TFs into four cross-species categories **(Figure 4i)**. Forty-two TFs showed concordant disease-associated activity shifts in both species, 102 were human-specific, 34 were mouse-specific, and 33 were significant in both species but changed in opposite directions. The mouse-specific and human-specific disease-associated networks contained 2,774 and 5,388 edges, respectively, with representative TF–target relationships shown in **Figure 4j** and complete records provided in Supplementary Table S5. Among the shared same-direction TFs, *E2F1* showed positive disease-associated activity shifts and *GATA3* showed negative activity shifts in both species **(Figure 4k)**. The concordant reduction in *GATA3* was consistent with its previously reported tissue-level downregulation in human psoriatic lesions and IMQ-treated mouse skin^48^. Several TFs showed opposing activity shifts: *SPI1*, *ETS1*, and *MYC* were positively associated with disease in mouse but negatively associated with disease in human, whereas *NFKB1* showed the opposite pattern. *STAT1* was identified as a human-specific disease-associated regulator **(Figure 4k)**. In addition, the top TFs regulating the main cell type populations differ between human and mouse, with Myc being the only shared dominant regulon **(Supplementary Figures S6,7)**. Collectively, these analyses showed that the IMQ mouse immune response differs substantially from the human response, with only a limited set of conserved features such as TNF signaling and concordant *E2F1* /*GATA3* activity changes **(Supplementary Figure S5)**.

### 2.5 A sparse single-cell gene panel diagnoses psoriasis with high accuracy and further reveals species-specific molecular signatures

Based on scSAID data, we further developed a machine-learning-based algorithm, psoSpotter, for psoriasis biomarker selection and applied it independently to the human psoriasis and mouse IMQ-induced psoriasiform inflammation datasets. The psoSpotter model training first restricted each species to the relevant binary disease–control contrast, ensuring that feature selection was driven by psoriasis-associated transcriptional differences rather than unrelated sample heterogeneity. Age and sex were retained as covariates to reduce the contribution of demographic or sample-composition effects to the classifier. The modeling framework consisted of four sequential feature-selection and validation steps. First, genes were ranked using a supervised disease-effect score, which quantified the standardized separation between disease and control cells. This step identified genes with strong diagnostic relevance while allowing both psoriasis-high and psoriasis-low features to contribute. Second, correlation-based redundancy pruning was used to remove highly co-expressed features; among genes with strong pairwise correlation, the gene with the larger disease-effect score was retained. This reduced the dominance of near-equivalent marker genes and increased the diversity of the candidate feature space. Third, ElasticNet-regularized logistic regression^49^ was applied across repeated subsampled training partitions as a stability-selection procedure^50^. Genes were ranked by their selection frequency across repeats, and the 50 most stable genes were retained as the final diagnostic panel in each species. The 50-gene size was chosen to balance interpretability and diagnostic performance, yielding a compact panel suitable for downstream biological interpretation and cross-species comparison. Finally, the selected 50-gene panel was evaluated in a logistic-regression classifier with age and sex covariates on held-out test cells **(Figure 5)**.

**Figure 5.**
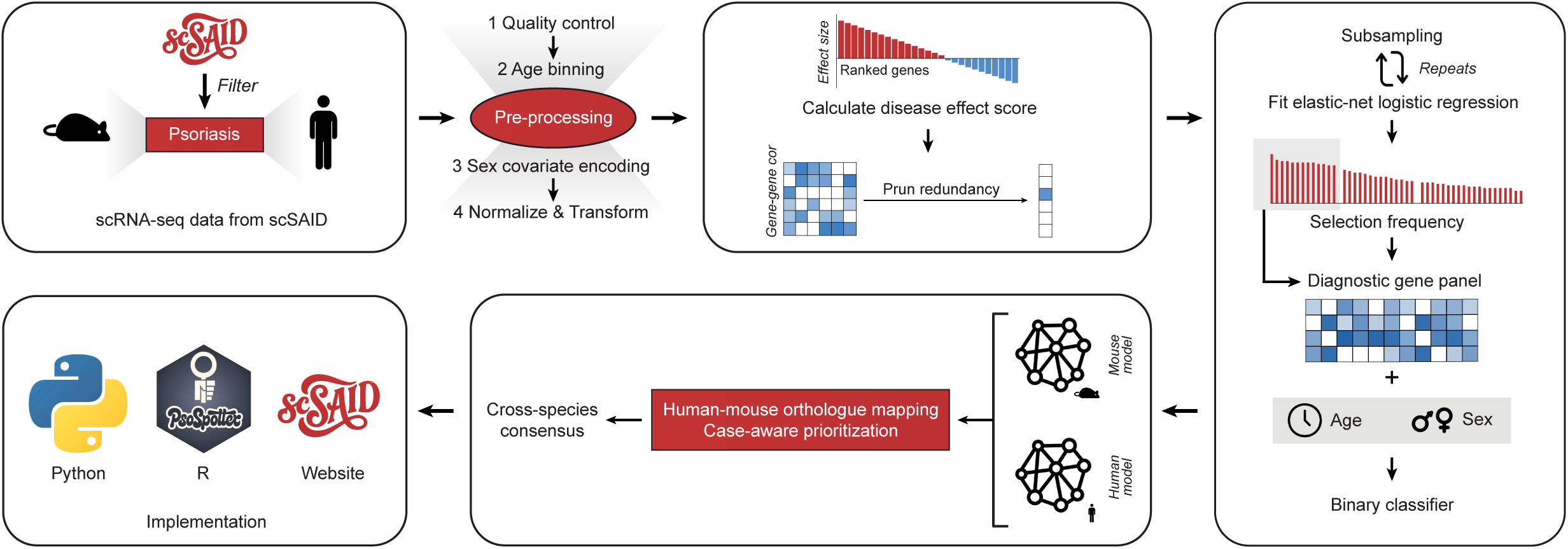
Overview of the psoSpotter modeling framework for cross-species psoriasis gene panel identification. scSAID-derived single-cell RNA-seq data were filtered to define species-specific binary diagnostic tasks, including healthy versus psoriasis cells in the human atlas and healthy versus IMQ-induced psoriasiform cells in the mouse atlas. The preprocessing workflow included quality control, age-bin assignment, sex-covariate encoding, library-size normalization and expression transformation before model training. Supervised feature filtering used a disease-effect score to rank genes by disease–control separation, followed by correlation-based redundancy pruning to remove highly co-expressed genes. Repeated subsampling and ElasticNet-regularized logistic regression were used for stability selection; genes were ranked by selection frequency and retained as a compact diagnostic gene panel. The selected diagnostic panel was combined with age and sex covariates in a binary classifier for psoriasis-state prediction. Human and mouse model outputs were integrated through ortholog mapping and case-aware prioritization to identify cross-species conserved and human-enriched psoriasis-associated candidate genes. The modeling workflow was implemented as a reproducible Python package, with additional implementations in R and on the scSAID website.

To facilitate reuse of the modeling framework, we implemented the algorithm as Python and R packages and as a scSAID website version (skin-scsaid.com/psospotter.jsp) for feature scoring, stability selection, diagnostic-panel construction and candidate-gene prioritization. This implementation separates disease-state modeling from psoriasis-specific labels, allowing users to provide new disease–control annotations and apply the same workflow to other single-cell disease atlases. Accordingly, psoSpotter provides a generalizable framework for building sparse disease classifiers and prioritizing disease-associated genes from annotated single-cell datasets.

By applying psoSpotter, we produced gene panels for both human and mouse. Since the final human and mouse diagnostic panels showed limited direct overlap, candidate target genes were prioritized from the broader stability-selection outputs rather than from the compact 50-gene panels alone. Genes with positive disease-associated coefficients in the human stability-selection output were retained as psoriasis-high candidates, and cross-species support was assessed using matched human–mouse orthologs. This procedure yielded 78 psoriasis-high candidates, including 17 cross-species conserved candidates with positive disease-associated coefficients in both species and 61 human-enriched candidates without matched positive mouse-model support **(Figure 6a–d)**. The leading candidate was *S100A8*, selected in 76% of human and 73% of mouse subsamples with positive disease coefficients in both models. Other conserved candidates included *S100A9*, *FOSB*, *IER2*, *RHOB* and *ATF3*, whereas the strongest human-specific candidates included *S100A7*, *PI3*, *HLA-DRB5* and *CCL27*, which we interpret as human-enriched psoriasis biology that the imiquimod model may not capture.

**Figure 6.**
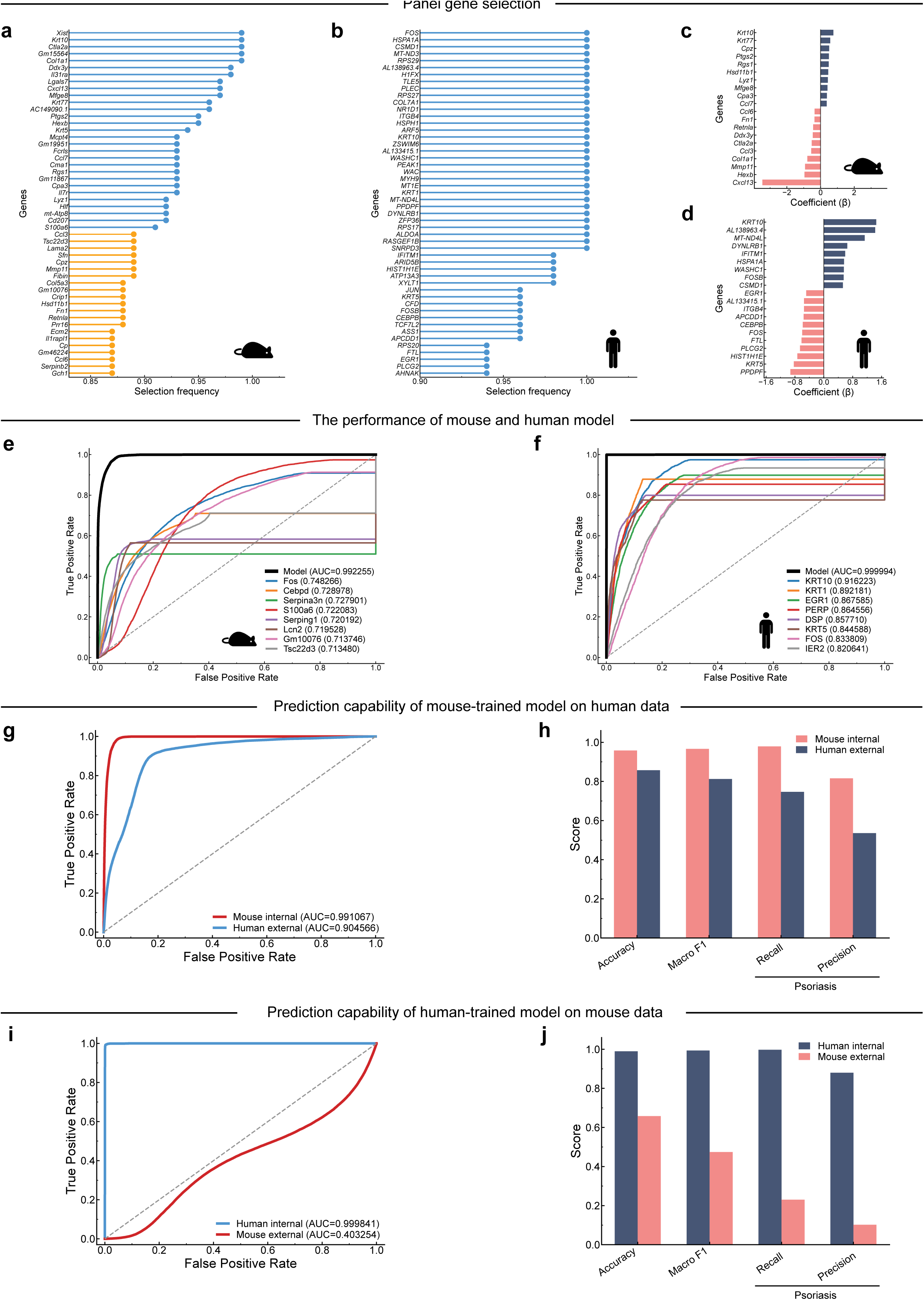
Performance of the cross-species psoriasis gene panel model in mouse and human skin. **(a,b)** Selection frequency of the top candidate genes identified by stability selection in the mouse **(a)** and human **(b)** datasets. **(c,d)** ElasticNet regression coefficients (*β*) of the final selected panel genes in the mouse **(c)** and human **(d)** models; positive values indicate association with psoriasis and negative values with healthy skin. **(e,f)** ROC curves for the mouse model **(e)** (AUC = 0.992) and human model **(f)** (AUC = 0.999), shown alongside individual-gene classifier performance for comparison. **(g)** Cross-species generalization of the mouse-trained model applied to human data (mouse internal AUC = 0.99; human external AUC = 0.91). **(h)** Bar chart summarizing accuracy, macro F1, psoriasis recall and psoriasis precision of the cross-species generalization described in **(g)**. **(i)** Cross-species generalization of the human-trained model applied to mouse data (human internal AUC = 1.00; mouse external AUC = 0.40). **(j)** Bar chart summarizing accuracy, macro F1, psoriasis recall and psoriasis precision of the cross-species generalization described in **(i)**.

The human classifier model reached near-perfect performance on held-out cells, with an accuracy of 0.998, an area under the receiver operating characteristic (ROC) curve (AUC) of 0.999, and macro and weighted F1 scores of 0.991 and 0.998 **(Figure 6f)**; it retained both sensitivity for psoriatic cells (recall 1.000, precision 0.968, F1 0.984) and specificity for healthy cells (recall 0.997, precision 1.000, F1 0.999). The mouse classifier model reached an AUC of 0.992, far above a label-shuffled control (AUC 0.47). In both species, the panel substantially outperformed the best individual genes, which reached an AUC of only approximately 0.92 in human (led by *KRT10* and *KRT5*) and approximately 0.75 in mouse (led by *Fos* and *Serpina3n*; **Figure 6e,f**).

Classification was therefore driven by a coordinated transcriptional signature rather than any single marker. Transfer of the panel between species was directional **(Figure 6g–j)**. The ortholog-restricted human-trained model retained perfect internal performance in human cells (AUC 1.00) but transferred poorly to the mouse dataset (AUC 0.40; **Figure 6i,j**), indicating that the human-derived diagnostic ranking was not preserved in the mouse IMQ model. In contrast, the reciprocal mouse-trained model retained high internal performance (AUC 0.99) and transferred effectively to human psoriasis (AUC 0.91; **Figure 6g,h**). This asymmetry indicates that the mouse model captures a conserved core of the human psoriasis state, while human psoriasis additionally engages disease-associated pathways that the mouse model does not fully reproduce.

### 2.6 Cross-platform in silico perturbation identifies four concordant candidates

To assess the predicted disease-reversal potential of the 78 psoSpotter-prioritized psoriasis genes, we compared Geneformer-simulated token deletion with gene-knockdown connectivity scoring against the LINCS L1000 resource^51,52^. Fifteen candidates had eligible Geneformer results (detection number *≥* 200 cells) and matched consensus shRNA signatures **(Figure 7a)**. Four genes—*RHOB*, *PPIA*, *PRKAR2B* and *HLA-DRB5* —showed concordant health-directed predictions and were classified as candidates with dual in silico support, whereas the remaining 11 candidates showed discordant directions **(Figure 7b,c)**. Among the four concordant candidates, *RHOB* had the most negative median XSum, indicating the strongest predicted LINCS knockdown reversal, whereas *PPIA* had the largest positive Geneformer Shift to goal end **(Figure 7d)**. Complete candidate-level results are provided in Supplementary Table S6, and workflow counts are provided in Supplementary Table S7.

**Figure 7.**
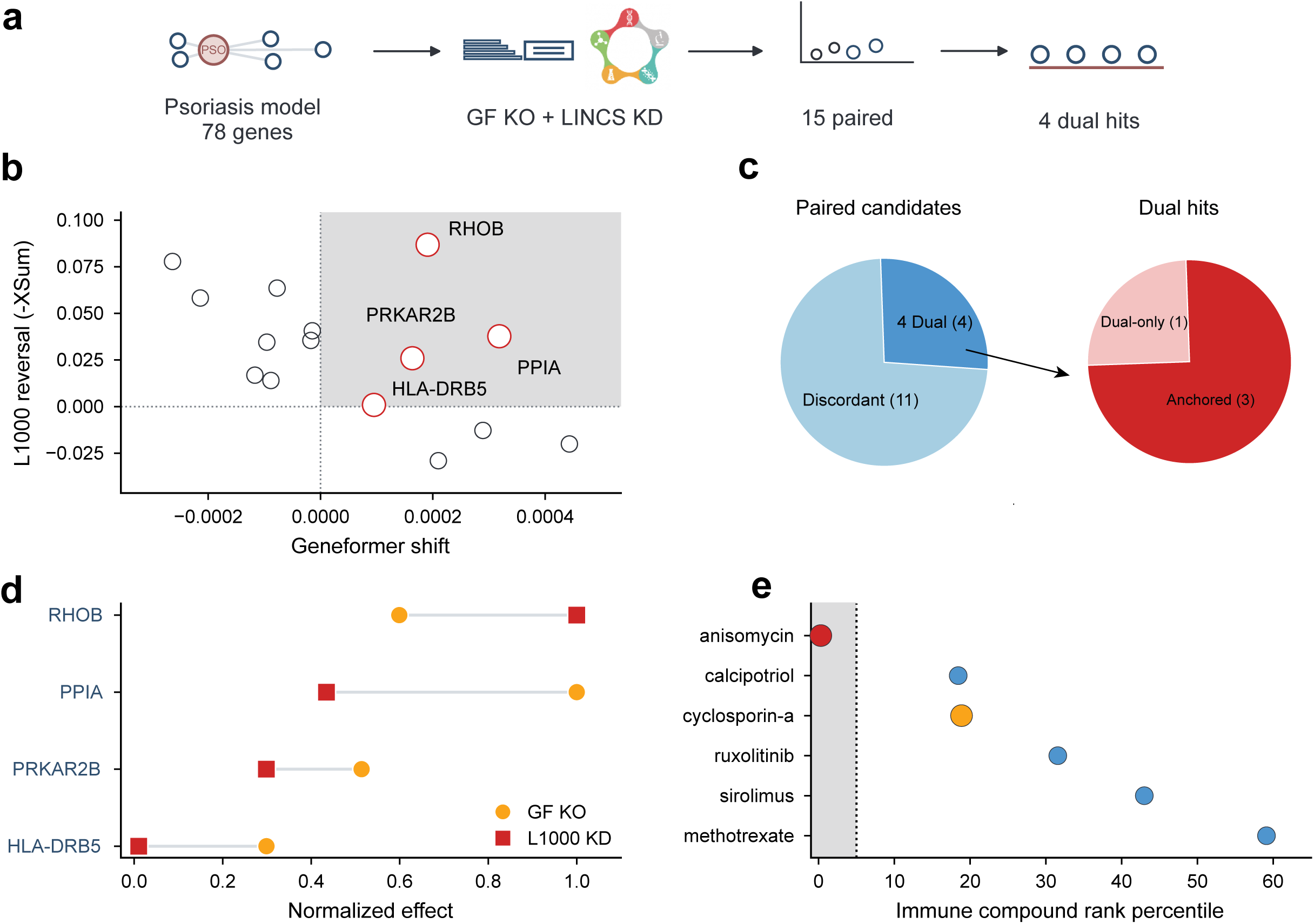
Cross-platform in silico perturbation analysis identifies concordant psoriasis candidates. **(a)** Schematic overview of the candidate-prioritization workflow. The 78 genes prioritized by the psoriasis classification models were evaluated using Geneformer token deletion and LINCS L1000 consensus shRNA signatures. Fifteen genes had eligible matched results, of which four satisfied the cross-platform concordance criterion. **(b)** Comparison of Geneformer Shift to goal end and LINCS reversal scores for the 15 paired candidates. Dotted lines indicate zero effect. The shaded upper-right quadrant represents Shift to goal end *>* 0 and median XSum *<* 0. **(c)** Classification of the 15 paired candidates into four candidates with concordant dual in silico support and 11 candidates with discordant directions (left). Of the four concordant candidates, *RHOB*, *PPIA* and *PRKAR2B* had additional mechanistic context, whereas *HLA-DRB5* was supported primarily by the cross-platform predictions (right). **(d)** Paired comparison of perturbation effect sizes for the four concordant candidates. Geneformer shifts and LINCS reversal scores were independently normalized to their respective maxima. **(e)** LINCS connectivity positions of selected compounds within the ranking of all 19,811 per-turbagens. Compounds were ranked by ascending median XSum, and the plotted reversal value is *−*median XSum. The dotted line marks the top 5%.

*PPIA* encodes cyclophilin A, the intracellular binding partner of cyclosporin^53^. The cyclosporin–cyclophilin complex inhibits calcineurin signalling^54^, a well-established immunosuppressive mechanism relevant to psoriasis^6^. Thus, the prioritization of *PPIA* identified a mechanistically interpretable candidate linked to cyclosporin pharmacology. Alongside *PPIA*, *RHOB* and *PRKAR2B* were retained as mechanism-anchored candidates, whereas *HLA-DRB5* was retained as a dual-only candidate because its psoriasis-specific mechanistic support was less direct. Compound connectivity did not identify a clear repurposing candidate. Cyclosporin A ranked 3,737th among 19,811 compounds (18.9th percentile), outside the top 5% of predicted reversers **(Figure 7e)**. Anisomycin ranked within the top 0.3%, but its ribotoxic-stress response and experimentally observed cutaneous psoriasiform effects^55,56^ make it an implausible psoriasis repurposing candidate despite its connectivity rank. The complete compound connectivity ranking is provided in Supplementary Table S8.

## 3 Methods

### 3.1 Data collection and curation

Raw FASTQ files generated using 10x Genomics droplet-based single-cell RNA-sequencing were downloaded from GEO or GSA. FASTQ files from each GSM accession were processed independently using cellranger count in Cell Ranger ^57^ v7.0.1, with the 10x Genomics refdata-gex-GRCh38-2020-A reference for human datasets or refdata-gex-mm10-2020-A for mouse datasets. Each dataset was annotated with species, condition, sex, age (or postnatal day for mouse), anatomical site, and per-sample cell count using the original metadata, then exported as a standardized AnnData object for downstream processing.

Each sample was imported into Scanpy (v1.11.4)^58^ as an AnnData object, and gene names were made unique. Mitochondrial, ribosomal, and hemoglobin genes were annotated, and per-cell quality-control metrics were calculated. Cells were retained if total counts, log-transformed total counts, and detected gene numbers fell between the 1st and 99th percentiles within each sample, and if mitochondrial, ribosomal, and hemoglobin transcript fractions were below 15%, 20%, and 5%, respectively. Putative doublets were identified with Scrublet^59^ and removed. The filtered count matrix was preserved in adata.layers[”counts”], followed by library-size normalization using the default sc.pp.normalize total setting and log1p transformation.

### 3.2 Integration

Quality-controlled samples were concatenated separately for each species using anndata.concat with an outer join, and batch groups containing fewer than 350 cells were excluded. To restrict model fitting to informative transcriptional features, highly variable genes (HVGs) were reselected from the raw-count layer using the seurat v3 procedure implemented in Scanpy. HVG selection was performed with batch as the stratification variable, and the top 8,000 genes were retained for each species. We adopted this feature-selection strategy on the basis of systematic atlas-level benchmarking, which has shown that restricting input to HVGs generally improves integration performance over the full gene set, both in overall batch-effect removal and in the conservation of biological variation^60^.

To obtain a batch-aware species-specific latent representation, species-specific integration was then performed using scVI (v1.3.3)^13^. Models were fitted to the raw counts stored in the counts layer, with batch specified as the batch key, GSE included as a categorical covariate, and the mitochondrial transcript fraction included as a continuous covariate. Each model comprised two hidden layers and a 30-dimensional latent space, with a negative-binomial gene-expression likelihood and a dropout rate of 0.1. Models were trained for up to 600 epochs with early stopping. The resulting latent representation was stored in X scVI and used to construct the integrated nearest-neighbor graph and UMAP embedding. Leiden clustering was subsequently evaluated across resolutions ranging from 0.05 to 1.0.

### 3.3 Cell-type annotation

Cell type identities were assigned by combining differential-expression analysis with manual review of canonical lineage markers (Supplementary Table S1). Prior to annotation, clusters containing fewer than 10 cells at any Leiden resolution from 0.05 to 1.0 were removed to exclude unstable low-abundance partitions. In the finalized workflow, Scanpy rank genes groups with the Wilcoxon test was applied to each Leiden resolution using the log1p layer. Marker genes were filtered by adjusted *P <* 0.05, log_2_ fold change *>* 0.5, and detection fraction *>* 0.1, and the top marker genes for each cluster were exported for manual inspection. Final labels were assigned at the resolution that produced the most biologically interpretable partition, including leiden scVI 0 8 for human after removal of cluster 4 and leiden scVI 0 7 for mouse after removal of cluster 10. Curated annotations were stored as fine-grained and broad labels in Fine Map and Gross Map.

### 3.4 Perturbation scoring

Cellular compartments most strongly perturbed by psoriasis were prioritized in each species using Augur^46^ via pertpy (v0.10.0)^61^. Disease contrasts were Healthy versus psoriasis for human and Healthy versus IMQ-induced psoriasis for mouse. Both Gross Map and Fine Map annotation levels were tested. Cell types with fewer than 50 cells per condition were excluded prior to scoring. Augur trains a random-forest classifier per cell type and reports a cell-type AUC that quantifies the magnitude of disease-associated transcriptomic shift. Default permutation-based significance, subsampling, and class-balancing were applied.

### 3.5 Differential expression and pathway enrichment

Cross-species comparisons were performed using human psoriasis datasets (SAID171, SAID173, SAID175, SAID176, SAID178 and SAID179) and mouse imiquimod-induced psoriasis-like datasets (SAID039, SAID092, SAID093, SAID110 and SAID111). Within each species, disease samples were compared with all available healthy controls. Pseudobulk count matrices were generated separately for each cell type and biological replicate using the aggregation utilities implemented in the decoupler package (v.2.1.1) ^62^. Differential expression analysis was subsequently performed with PyDESeq2 (v.0.5.2)^63^, which fits a negative-binomial generalized linear model to pseudobulk count data. The batch variable was used as the biological-replicate identifier, whereas condition was specified as the contrast variable. The same disease–control contrasts as those used in the Augur analysis were applied. Genes with adjusted *P <* 0.05 (Benjamini–Hochberg procedure) and absolute log_2_ fold change *>* 0.5 were considered differentially expressed.

For cross-species comparison, mouse gene symbols were mapped to human orthologs through Ensembl BioMart^64,65^ using pybiomart (v.0.2.0; https://github.com/jrderuiter/pybiomart). Only one-to-one ortholog relationships were retained for direct human–mouse comparisons. Significant DEGs were classified as shared when their one-to-one ortholog was also significantly differentially expressed in the other species, species-specific when a one-to-one ortholog was available but was not significantly differentially expressed in the other species, or no-ortholog when no one-to-one ortholog could be identified. Genes in the no-ortholog category were excluded from direct cross-species overlap calculations.

ORA was performed with GSEApy (v.1.1.11)^14^ using gseapy.enrichr and locally downloaded GMT files from the human MSigDB v2026.1.Hs and mouse MSigDB v2026.1.Mm releases. Mouse-specific gene lists were tested against the MH Hallmark, M5 ontology, and M8 cell-type-signature collections, whereas human-specific gene lists were tested against the corresponding H Hallmark, C5 ontology, and C8 cell-type-signature collections. Shared genes were represented by their human ortholog symbols and analyzed against the human H, C5, and C8 collections. Enrichment significance was assessed using Fisher’s exact test, followed by Benjamini–Hochberg correction for multiple testing. Terms with an adjusted *P <* 0.05 were considered significant.

### 3.6 Cell-cell communication inference

Cell–cell communication was inferred separately for each (species, condition) combination using CellChat (v2.2.0)^66^ with the species-matched CellChatDB. The atlas was restricted to comparable contexts (whole skin samples in the human atlas, all samples in the mouse atlas) and split by condition. computeCommunProb was run with type = “triMean” and trim = 0.1, followed by filterCommunication(min.cells = 10), computeCommunProbPathway, and aggregateNet. Per-pathway signaling-strength heatmaps and incoming and outgoing role plots for the immune compartment subset were generated through netVisual heatmap and netAnalysis signalingRole heatmap.

### 3.7 Gene regulatory network inference

For human psoriasis and the mouse IMQ-induced psoriasis-like model, sample-level gene regulatory networks were inferred for the immune compartment using SCORPION (v.1.3.2)^20^. DoRothEA interactions with confidence levels A–C ^67^ and TF-TF protein interactions from STRING v12.0 were used as regulatory priors. Networks were constructed for sample–condition groups containing at least 30 cells, using 25 principal components, a smoothing parameter of *γ* = 10, and Pearson correlation.

Disease-associated TF–target edge-weight changes were assessed by comparing disease and healthy sample-level networks with SCORPION’s testEdges function, followed by Benjamini–Hochberg correction. TF-centered summaries were calculated as the mean disease-minus-healthy edge-weight change across the targets associated with each TF. For cross-species comparison, mouse genes were mapped to one-to-one human orthologs using the R package biomaRt v2.62.1. Orthologous TFs and TF–target edges were classified as concordant or divergent according to the direction of their disease-associated changes. Conserved edges were required to show the same direction and an adjusted *P <* 0.1 in both species, whereas conserved TF candidates were required to show concordant mean edge-weight changes and at least one supporting edge with an adjusted *P <* 0.1 in each species.

### 3.8 psoSpotter framework

#### 3.8.1 Feature filtering and stability selection

For the single-cell diagnostic model, each species’ atlas was converted into a stripped AnnData object retaining only X, obs, and var, then exported as a Zarr store^68^ after filtering. Filtering required parseable age annotations, gene detection in *≥* 1% of age-eligible cells, library-size normalization to 10,000 counts per cell, and log1p transformation using Scanpy. Age was treated as a required covariate. Human ages were assigned to four bins (0–20, 20–40, 40–60, and *>* 60 years) and mouse ages to four bins (0–6 weeks, 6 weeks to 6 months, 6 months to 1 year, and *>* 1 year). Sex was encoded as 1 (male), 0 (female), or 0.5 (ambiguous or missing). For each gene, a supervised effect score was computed as

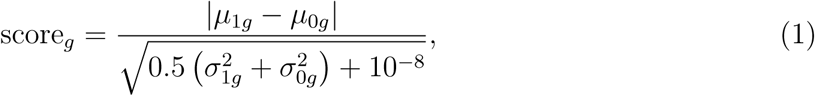

where *µ_kg_* and 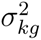 are the mean and variance of expression for gene *g* in disease (*k* = 1) and healthy (*k* = 0) cells. Genes with absolute pairwise Pearson correlation *≥* 0.90 in the training set (estimated on up to 15,000 randomly sampled cells with block-wise sufficient statistics) were pruned, retaining the higher-scoring gene per redundant pair. Stability selection was then applied^50^: 30-100 repeats depending on the task, 75% of training groups were subsampled, and an ElasticNet logistic regression was fitted with class-balanced weights, l1 ratio = 0.9, and C tuned in [10^−5^, 10^−2^] to target approximately 250 non-zero coefficients. Genes were ranked by selection frequency, and the top 50 were retained as the final diagnostic panel.

#### 3.8.2 Training, evaluation, and cross-species transfer

The final classifier was a logistic-regression model fitted with the SAGA solver^69^ on the 50-gene panel together with age-bin and sex covariates using scikit-learn^70^:

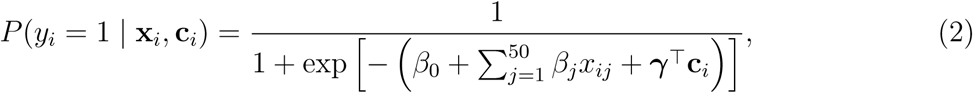

where *y_i_*= 1 denotes psoriasis (or IMQ-induced psoriasis in the mouse model), *x_ij_* is the *z*-scored expression value of gene *j* in cell *i* clipped to [*−*5, 5], and **c***_i_* is the covariate vector. Performance was evaluated using an 80:20 stratified group-held-out split with a fixed random seed; splitting was performed at the donor group level to prevent cells from the same biological group from being assigned to both partitions. Feature selection and model fitting were restricted to the training partition. For the human model, the panel was additionally assessed with stratified group *k*-fold cross-validation. For the mouse model, a train-label shuffle was used as a negative control. Metrics included accuracy, ROC-AUC, precision, recall, macro F1, and weighted F1. For cross-species transfer, the analysis was restricted to one-to-one orthologs with mitochondrial and ribosomal genes excluded. Age was harmonized across species into four shared developmental bins: juvenile (*<* 18 years in human, *<* 6 weeks in mouse), young adult (18–40 years, 6 weeks to 6 months), adult (40–65 years, 6–12 months), and aged (*≥* 65 years, *≥* 1 year). A model trained in one species was projected onto the matched orthologous genes in the other and re-evaluated. Candidate target genes were prioritised from the broader stability-selection outputs rather than from the compact final 50-gene diagnostic panels alone. Genes with positive mean disease-associated coefficients in the human stability-selection output were retained as psoriasis-high candidates. Cross-species conserved candidates were defined as genes with matched human–mouse orthologues and positive disease-associated coefficients in both species. Human-enriched candidates were defined as human psoriasis-high genes without matched positive mouse-model support.

### 3.9 Geneformer zero-shot in silico gene deletion

The 78 model-prioritized candidates were mapped to Ensembl identifiers and evaluated against the Geneformer V2 gc104M vocabulary. Sixty-seven candidates, including 17 with cross-species support, were retained for analysis. Geneformer (v.0.1.0)^51,71^ with the pretrained Geneformer-V2-104M model was applied to raw-count profiles from the immune compartment. Cells were tokenized as rank-ordered gene sequences with a maximum length of 4,096 genes.

Each candidate was evaluated in zero-shot mode by deleting its token from psoriasis-cell sequences. Perturbation effects were summarized using Geneformer’s goal state shift mode. The resulting Shift to goal end statistic quantifies the cosine shift of perturbed cell embeddings from the psoriasis state towards the healthy-state representation. Up to 2,000 cells were evaluated per candidate, and candidates with an effective sample size below 200 cells were excluded. Analyses were performed in FP32 on a single NVIDIA V100 GPU.

### 3.10 LINCS L1000 perturbation analysis

To complement the Geneformer predictions with experimentally derived perturbation profiles, we used the LINCS L1000 pilot-phase dataset deposited under GEO accession GSE92742^52^. We analysed Level 5 signatures, in which replicate profiles are combined into differential-expression signatures representing the transcriptional response to genetic or compound perturbation. The query comprised up to 150 significantly upregulated and 150 significantly downregulated genes, selected at adjusted *P <* 0.05 and ranked by log_2_ fold change.

For each L1000 signature *s*, connectivity was calculated using a matched-gene-normalized implementation of XSum^72^:

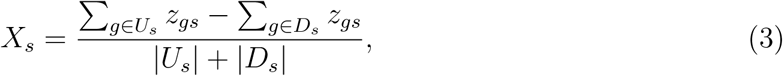

where *U_s_* and *D_s_* are the upregulated and downregulated query genes represented in signature *s*, and *z_gs_* is the corresponding Level 5 value. Scores were summarized by the median across all available signatures for each perturbagen, pooling cell lines, doses and exposure times. Negative median XSum values indicate predicted reversal of the psoriasis-associated expression programme. For visualization in **Figure 7**, the sign-inverted value (*−*median XSum) was plotted so that stronger predicted reversal appears in the positive direction.

Consensus shRNA signatures (trt sh.cgs) were used to assess candidate-gene knockdown. Candidates with Geneformer N Detections*≥* 200 and an available consensus-shRNA signature were considered to have concordant dual in silico support when Shift to goal end *>* 0 and the median XSum was negative. For compound prioritization, all available compound signatures (trt cp) were evaluated using the same psoriasis query and ranked from the most negative to the most positive median XSum. Mechanism-of-action and target annotations for selected compounds were obtained from the Drug Repurposing Hub^73^.

### 3.11 Web platform server construction

scSAID runs on an Ubuntu 24.04.3 LTS x86-64 machine equipped with a 4-vCPU Intel Xeon Platinum processor and 16 GB of RAM. Its backend is built chiefly in Java on the Java Servlet/JSP framework, running on Apache Tomcat 9.0.115 that sits behind an Nginx reverse proxy. User-specific computational jobs are kept separate through server-side HTTP sessions, whereas the public portal is open and requires no accounts. On the client side, the interface is assembled from JSP-rendered HTML5, CSS3, and JavaScript, combining a custom CSS component system with jQuery and DataTables to drive interface behavior and tabular displays, alongside Plotly.js and ECharts for interactive data visualization. The frontend and backend communicate via asynchronous HTTPS requests issued through the Fetch API and jQuery AJAX, with JSON serving as the main data-exchange format.

## 4 Discussion

In this study, we developed scSAID as a disease-oriented, cross-species single-cell resource for skin biology and skin disease. scSAID integrates 10x skin single-cell RNA-sequencing datasets across species, disease states, tissue sites, cell types and experimental conditions, transforming fragmented public datasets into a framework for asking how skin disease programs are conserved, remodeled, or lost across biological contexts.

The major biological insight of this study using scSAID emerges from the integrative cross-species analysis. Both the human and imiquimod-induced mouse immune compartments were strongly disease-perturbed, supporting continued use of the imiquimod model for selected inflammatory questions. However, transcriptional, signaling, and regulatory comparisons showed that shared perturbation does not imply biological equivalence. Human psoriasis was more strongly associated with leukocyte and T-cell migration, NF-*κ*B-related activity, and STAT1-associated interferon circuitry, whereas the imiquimod model showed stronger associations with acute myeloid and proliferative programs and with epithelial-junction or keratinization-associated features. This pattern is consistent with prior work showing that imiquimod-induced dermatitis activates psoriasis-related pathways while displaying strain dependence, acute kinetics, and a broad core skin-inflammation signature rather than a psoriasis-specific chronic program^12,74^. The imiquimod model is therefore not simply valid or invalid, but modularly valid. This distinction is important because heavy reliance on the IMQ model may have constrained mechanistic inference for human psoriasis by conflating model-specific inflammation with disease-general biology. Previous studies have raised this concern^75^, but have largely criticized the model as a whole rather than defining which components remain informative. Our analysis addresses this gap by separating conserved inflammatory modules from species-specific or discordant programs that should not be directly extrapolated to human disease.

Classifier transfer with psoSpotter using scSAID data provides an independent quantitative test of this modular view. High internal performance of both classifiers indicates that psoriasis-associated cell states are encoded by coordinated transcriptional programs. The transfer was asymmetric: the mouse-trained model generalized to human psoriasis, whereas the human-trained model did not generalize to the imiquimod model. This supports a directional model of conservation in which the mouse captures a subset of the human psoriasis state, while human lesions contain additional chronic, adaptive, and tissue-contextual programs. Treatment-response single-cell studies are consistent with our results, showing that pathogenic tissue-resident T-cell states and inflammatory niches are remodeled during IL-23 blockade^76,77^. In addition, the psoSpotter algorithm supports selection of a minimal set of psoriasis marker genes suitable for age- and sex-aware testing, which is desirable in a clinical setting, and can also be applied to other diseases to select marker genes. Clinically, the main value of the diagnostic model is that it reduces a high-dimensional single-cell transcriptomic profile to a minimal testing gene set. Rather than requiring genome-wide expression measurement for every application, the stability-selected panel identifies a compact group of genes that preserves diagnostic information and can guide the design of targeted assays. This sparse-panel strategy may support more practical downstream validation, including focused qPCR or spatial/in situ assays, while also providing a ranked set of candidate genes for perturbation-based target evaluation.

scSAID is updated monthly as new skin single-cell datasets become available, ensuring that users have access to recent data. The analytical scope also continues to expand, most recently through functional module analysis using MAPA (https://github.com/jaspershen-lab/mapa), which links related enrichment terms into interpretable modules. We also recognize the pressure that computationally intensive analyses place on the server. Analysis-heavy queries such as cross-sample differential expression are therefore handled through a job queue that limits the number of concurrent submissions, ensuring that computationally demanding analyses remain available without degrading interactive performance. A backup server is planned to preserve access during maintenance. Future versions of scSAID will incorporate single-cell chromatin-accessibility and spatial-transcriptomic data, connecting the transcriptional states described here to regulatory elements and tissue architecture.

In summary, we developed scSAID as an integrated, queryable resource for exploring skin cell states, disease signatures and intercellular communication across species. We demonstrated its utility by reanalyzing a published dataset with advanced analysis features, performing an integrative analysis of species-specific psoriasis and generating new insights. With monthly updates and continued expansion, scSAID converts a decade of fragmented public skin datasets into a living foundation for cross-species disease research.

## 5 Acknowledgment

We thank National Key Research and Development Program of China (2021YFA1101100) for sponsoring this work. We would also like to acknowledge all members of the Chaochen Wang group and Ying Xiao group for their kind help in providing advice on the website and database with special thanks to Yanzhi Bian for his valuable advice on the manuscript.

## 6 Data availability

All raw sequencing data analyzed in this study were obtained from publicly available datasets deposited in GEO and GSA. The complete list of GEO and GSA accession numbers used to construct the scSAID resource is provided in Supplementary Table S9. The scSAID website is freely accessible at skin-scsaid.com. The final processed integrated h5ad objects are available upon publication.

## 7 Code availability

Source code for the scSAID database, web platform, and analysis pipelines is available upon publication. The R version of psoSpotter is available at https://github.com/EthanShenx/ psoSpotter-r; a Python version is available at https://github.com/EthanShenx/psoSpotter-py.

## 8 Author contribution

C.W. and Y.X. conceived the study. C.W., Y.X., Y.R., Y.S., and Y.H. contributed to dataset collection. Y.R. and Y.S. contributed to analysis. Y.S., Y.R., Y.D., and Y.H. developed and implemented the website. Y.S. maintains it. L.J. implemented the diagnostic model, with the help of Y.R. and Y.S. Y.S., Y.R., and L.J. interpreted the results. Y.R., Y.S., and L.J. made the figures and drafted the manuscript. C.W., Y.X, Y.S., Y.R. and L.J. revised the final manuscript.

## 9 Supplementary information

Supplementary information is available for this paper in separate Supplementary Information files. All items are cited in the main text.

## 10 Declarations

- **Competing interests:** The authors declare no competing interests.
- **Consent for publication:** All authors consent for publication.

